# CTCF-anchored cohesin loops are a vertebrate-specific innovation

**DOI:** 10.64898/2026.11.16.767090

**Authors:** Histoshi Aihara, Takeo Narita, Elina Maskey, Thi Binh Nguyen Nguyen, Yoshiki Higashijima, Yoko Yamaguchi, Nils Krietenstein, Chunaram Choudhary

**Affiliations:** Proteomics Program, The Novo Nordisk Foundation Center for Protein Research, Department of Cellular and Molecular Medicine, Faculty of Health and Medical Sciences, University of Copenhagen, Blegdamsvej 3B, 2200 Copenhagen, Denmark; Institute for Tenure Track Promotion, University of Miyazaki, Miyazaki, Japan; Institute of Agricultural and Life Sciences, Academic Assembly, Shimane University, Shimane, Japan; Faculty of Life and Environmentaol Sciences, Shimane University, Shimane, Japan; Center for Epigenetic Cell Memory, Danish Cancer Society, Copenhagen, 2100, Denmark

## Abstract

Animal genomes are organized into chromatin loops and topologically associating domains. Cohesin drives loop extrusion, and the architectural factor CTCF halts loop-extruding cohesin. CTCF-anchored loops facilitate interactions between distal cis-regulatory elements and regulate gene expression. However, when CTCF acquired its cohesin loop-halting function remains unclear. We performed interspecies domain-swap experiments, replacing the cohesin-interacting N-terminus of mouse CTCF with orthologous sequences from the fruit fly *Drosophila* (arthropod), *Branchiostoma* (chordate), and the inshore hagfish (jawless vertebrate). The hagfish N-terminus halted cohesin, whereas the invertebrate CTCF N-termini did not. Consistently, Micro-C maps from hagfish tissues revealed stripes associated with convergently oriented CTCF sites, a characteristic signature of cohesin-loop halting. Notably, cohesin enrichment at promoters and enhancers was more pronounced in hagfish than in jawed vertebrates. Our findings indicate that CTCF-mediated cohesin loop anchoring most likely evolved in the common ancestor of extant vertebrates and was further strengthened in jawed vertebrates.

## Introduction

Animal genomes are organized into chromatin loops and topologically associating domains (TADs)^1,2^. Chromatin loops have been reported in both bilaterian and non-bilaterian animals^3^, but the mechanistic principles of genome folding have been elucidated mostly in bilaterians. Of the factors involved in three-dimensional (3D) genome folding, cohesin and the C2H2 zinc-finger protein CTCF (CCCTC-binding factor) are the best studied. Although CTCF is broadly present across bilaterians^4^, its functions differ in *Drosophila* and mammals. *Drosophila* CTCF is enriched mainly at insulator boundaries, which block enhancers from activating illegitimate promoters^5,6^. In mammals, by contrast, CTCF acts as a cohesin loop anchor^7,8^. In the prevailing model, cohesin extrudes chromatin loops until it encounters convergently oriented CTCF binding sites, which halts extrusion^9^.

This genome organization is crucial for gene regulation. Mammalian enhancers can activate promoters located many hundreds of kilobases away^10,11^. Nonetheless, enhancer activity decays with distance^12,13^, likely because stochastic interactions between enhancers and promoters become less frequent as the distance between them increases. Cohesin-mediated loop extrusion can bridge this gap by bringing distal enhancers and promoters into 3D proximity. By halting cohesin, CTCF can restrict enhancer interactions to promoters within the same loop domain and prevent enhancers from activating promoters in neighboring domains^9,14^. This CTCF–cohesin-dependent genome organization is therefore important for enhancer specificity, particularly because mammalian enhancers often lack strong intrinsic promoter specificity in their native chromatin environment^12^. Consistent with this role, inverting CTCF-binding motif orientation can disrupt enhancer–promoter interactions and lead to inappropriate gene activation^7,15^. Similarly, structural variants that alter three-dimensional genome organization can cause severe developmental defects^16,17^.

Although mammalian CTCF also regulates gene expression by functioning as a transcriptional activator and repressor^18–20^, its role in halting cohesin-mediated loop extrusion has drawn most attention^21^. This is because the way CTCF promotes mammalian genome organization is unique. CTCF remains the only architectural protein whose ability to halt cohesin-mediated loop extrusion has been demonstrated biochemically, structurally, and genetically. However, when this hallmark function of CTCF evolved remains an outstanding open question.

Here, by analyzing the evolutionary conservation of CTCF and its cohesin-interacting motifs, swapping CTCF N-terminal regions between species, and performing Micro-C analyses, we show that CTCF’s cohesin-loop-halting activity likely evolved in the last common ancestor of crown vertebrates. The findings suggest that arthropods and vertebrates independently co-opted CTCF for distinct roles in genome organization, and that CTCF-anchored cohesin loops are a vertebrate-specific innovation.

## Results

### Cohesin-interacting CTCF motifs are not universally conserved

To investigate the evolutionary origin of CTCF-anchored loops, we examined the conservation of CTCF itself and two motifs in its N-terminus implicated in cohesin-mediated loop blocking: YxF (mouse 226-228) and KTYQR (mouse 23-27)^22–24^. Consistent with previous observations^4^, CTCF orthologs are not detectable in nematodes or platyhelminthes (**Figure 1A**). These lineages also exhibit highly derived genomic and regulatory features. In nematodes, for example, a substantial proportion of genes are organized into operons^25^. Platyhelminthes also have a reduced complement of transcription factors and developmental regulators that are broadly conserved across other bilaterians, including homeobox and Wnt genes^26,27^.

**Figure 1:**
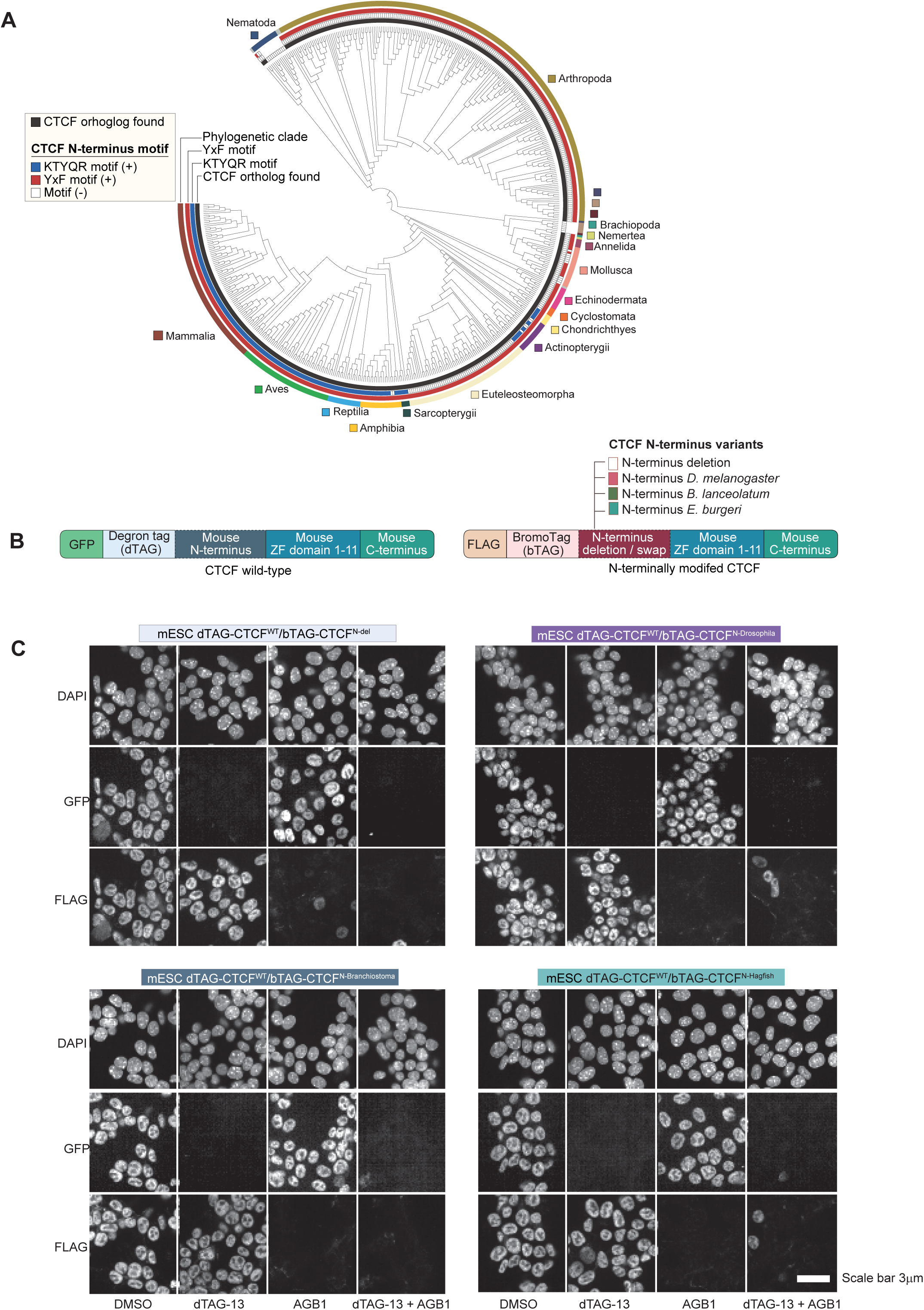
Evolutionary analysis of CTCF motif conservation and generation of CRISPR-edited cell models. **(A)** Phylogenetic tree showing conservation of CTCF and the cohesin-interacting motifs in CTCF across 527 bilaterians. The conservation of the two known cohesin-interacting motifs^22–24^, YxF and KTYQR, is indicated. Presence of the KTYQR and YxF motifs was determined from a multiple sequence alignment of CTCF N-terminal regions using MAFFT L-INS-i algorithm. For each sequence, the residues aligned to human CTCF positions 23–27 (KTYQR) and 226–228 (YDF) were examined. KTYQR was marked as present only when all five residues were identical to the human sequence. YxF was scored as present when the aligned residues matched [Y/F]-x-[Y/F] (x, any residue). All the species name and their motif annotation are listed in **Supplementary Table3**. **(B)** Schematic representation of degron-fused CTCF wild-type (CTCF^WT^) and N-terminally modified CTCF variants expressed in mouse ESC. In the deletion variant, the N-terminal region of mouse CTCF (amino acids 1–226) was removed. In the domain-swap variants, this region was replaced with the corresponding N-terminal sequence from the indicated species. The generation of cell lines expressing the modified CTCF variants in an allele-specific manner is described in **Supplementary Figure 1**. **(C)** Representative immunofluorescence images showing expression of the indicated CTCF variants in the corresponding cell lines. Cells were treated with dTAG-13 and AGB1 for 2 h before analysis. GFP staining indicates expression of CTCF^WT^, while FLAG immunostaining detects the N-terminally modified CTCF variants (n = 3 biological replicates).

The YxF motif is broadly conserved across bilaterians, although it is not universal. Notably, it appears to be absent from many molluscs. The KTYQR motif is even less conserved and is absent from most bilaterian groups, including arthropods and molluscs. Among vertebrates, KTYQR is restricted to jawed vertebrates; even in this group, it is apparently absent from euteleostomorph fishes. Together, these findings indicate that YxF is more broadly conserved than KTYQR, which appears to be restricted to a subset of jawed vertebrates.

### Generation of a cell model for assessing CTCF loop anchoring function

Mammalian CTCF has two distinct regulatory functions: it anchors cohesin-mediated loops and acts as a canonical transcription factor. These functions are mediated by distinct regions of the protein: the N-terminus is responsible for halting cohesin, whereas the C-terminus mediates transcriptional activation ^20,23,24,28^. CTCF-mediated transcriptional activation is essential for the expression of many housekeeping genes and for mammalian cell viability ^20^. To investigate the evolutionary origins of CTCF-anchored cohesin loops, we selectively deleted or swapped the N-terminus of mouse CTCF while preserving its DNA-binding and C-terminal domains of mouse. The mouse CTCF N-terminal sequences was swapped with the N-terminus of CTCF from three evolutionarily distant species: the arthropod *Drosophila melanogaster*, the cephalochordate *Branchiostoma lanceolatum*, and the cyclostome (jawless vertebrate) *Eptatretus burgeri* (inshore hagfish).

Using CRISPR, we engineered mouse stem cells to express wild-type and N-terminally modified CTCF from separate endogenous *Ctcf* alleles. Each protein was fused to a distinct degron tag, allowing selective and acute depletion (**Supplemental Figure 1A**). The wild-type allele was fused at its N-terminus to a dTAG, enabling selective depletion of wild-type CTCF. The second allele was fused to a BromoTag (bTAG). In this allele, we either deleted the N-terminal region (amino acids 1–265) or replaced the mouse N-terminus with the corresponding sequence from one of the three species. Notably, mice heterozygous for *Ctcf* are viable and phenotypically normal ^29^, suggesting that a single functional allele is sufficient to support normal development.

We generated four difference cell lines. The one expressed dTAG-fused wild-type CTCF and bTAG-fused CTCF lacking the N-terminal region (CTCF^WT^/CTCF^N-del^). The remaining three expressed dTAG-fused wild-type CTCF together with bTAG-fused chimeric CTCF proteins. Each chimeric protein retained the mouse zinc-finger domains (ZF1–11) and C-terminal region but contained the N-terminal region from either *Drosophila* (CTCF^WT^/CTCF^N-Drosophila^), or *branchostoma* (CTCF^WT^/CTCF^N-Branchiostoma^), or inshore hagfish (CTCF^WT^/CTCF^N-Hagfish^).

This allele-specific system offers three key advantages. First, both CTCF variants remain under endogenous regulatory control, minimizing differences in expression levels. Second, their distinct degron tags allow either variant to be depleted independently or in combination. Third, because the variants share identical mouse DNA-binding and C-terminal domains, any differences in cohesin regulation can be attributed to changes in the N-terminus. Moreover, because the N-terminally modified proteins retain the C-terminal transcriptional activation domain, cells can remain viable following selective depletion of either wild-type or N-terminally modified CTCF.

For all four cell lines, we confirmed CTCF expression, appropriate nuclear localization, and allele-specific degradation of the degron-tagged proteins following treatment with dTAG13 and AGB1, which induce degradation of dTAG- and bTAG-tagged proteins, respectively (**Figure 1B**). To confirm that the heterologous N-terminally swapped did not impair DNA binding, we performed ChIP-seq in CTCF^WT^/CTCF^N-Branchiostoma^. CTCF^N-Branchiostoma^ showed robust chromatin binding, which was abolished following AGB1 treatment (**Supplemental Figure 1B**).

### Invertebrate CTCF N-terminus fails to robustly enrich RAD21

To investigate the role of the CTCF N-terminus in cohesin retention, we treated all four cell lines with dTAT-13 and AGB1 (4 h), either individually or in combination, and examined genomic RAD21 occupancy by ChIP-seq. The four cell lines showed variable reductions in RAD21 enrichment following selective depletion of CTCF^WT^. However, in all four cell lines, including CTCF^WT^/CTCF^N-del^ cells, combined depletion of CTCF^WT^ and the N-terminally modified CTCF variant caused a greater reduction in RAD21 enrichment than depletion of either variant alone (**Supplementary Figure 2**). These results suggest that, although the CTCF N-terminus is important for robust RAD21 enrichment, N-terminally truncated CTCF retains some capacity to support cohesin enrichment. This basal RAD21 enrichment may be mediated by CTCF’s zinc-fingers.

We first examined changes in RAD21 occupancy following combined depletion of CTCF^WT^ and the N-terminally modified CTCF variant across all four cell lines. At CTCF-positive (bound by CTCF, CTCF^+^) peak regions, RAD21 occupancy was globally reduced. In contrast, RAD21 occupancy remained unaffected at a subset of CTCF-negative (not bound by CTCF, CTCF^−^) peak regions (**Figure 2A**).

**Figure 2:**
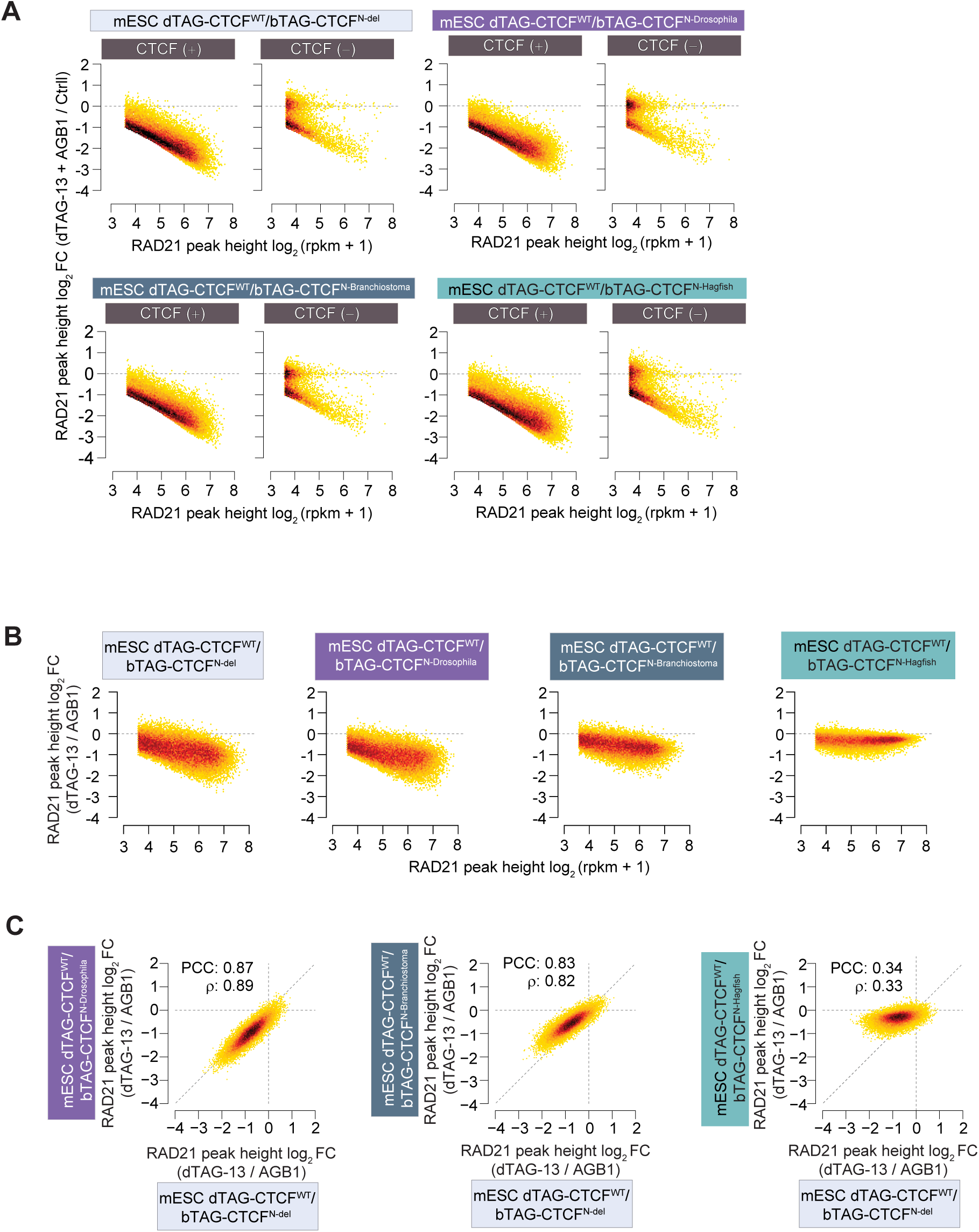
N-termini of jawed and jawless vertebrate CTCF support cohesin enrichment in mouse ESC. **(A)** RAD21 occupancy following simultaneous depletion of CTCF^WT^ and the indicated N-terminally modified CTCF variant. Cells were treated with dTAG-13 and AGB1 simultaneously for 4 h. Genome-wide RAD21 occupancy was measured by ChIP-seq. Shown are fold changes in RAD21 peak heights in CTCF-depleted cells relative to untreated control cells (n = 2 biological replicates). RAD21 occupancy is compared in CTCF^+^ and CTCF^−^ regions. **(B)** RAD21 occupancy following selective depletion of either CTCF^WT^ or the indicated N-terminally modified CTCF variants. Cells were treated with either dTAG-13 or AGB1 for 4 h to selectively deplete CTCF^WT^ and the N-terminally modified CTCF variants, respectively. Genome-wide RAD21 occupancy was measured by ChIP-seq. Shown are fold changes in RAD21 peak heights in the indicated treatment conditions (n = 2 biological replicates). **(C)** Correlation of RAD21 peak-height fold changes between the indicated cell lines following dTAG-13 or AGB1 treatment. In each cell line, RAD21 peak-height fold change was calculated by comparing dTAG-13-treated and AGB1-treated conditions.

Based on this observation, we focused on CTCF^+^ peak regions to assess changes in RAD21 enrichment following selective depletion of CTCF^WT^. In CTCF^WT^/CTCF^N-del^ cells, selective depletion of CTCF^WT^ resulted in a global reduction in RAD21 enrichment, confirming that CTCF^N-del^ cannot efficiently support cohesin enrichment (**Figure 2B**). A comparable reduction was observed in CTCF^WT^/CTCF^N-Drosophila^ cells after CTCF^WT^ depletion. Depletion of CTCF^WT^ also led to strong reduction in RAD21 in CTCF^WT^/CTCF^N-Branchiostoma^ cells, whereas CTCF^WT^/CTCF^N-Hagfish^ cells showed the smallest reduction.

To directly compare the ability of the different N-terminally modified CTCF variants to support cohesin enrichment, we examined the correlations in RAD21 enrichment changes across cell lines following depletion of CTCF^WT^, when only the modified CTCF variants remained expressed. RAD21 enrichment changes were strongly correlated between CTCF^N-del^ and CTCF^N-Drosophila^ cells (Pearson’s r = 0.87; Spearman’s ρ = 0.89; **Figure 2C**), indicating comparable effects on cohesin enrichment. Similarly, RAD21 enrichment changes in CTCF^N-Branchiostoma^ cells were strongly correlated with those in CTCF^N-del^ cells (Pearson’s r = 0.83; Spearman’s ρ = 0.82), although the reduction was slightly less pronounced in CTCF^N-Branchiostoma^ cells. In contrast, RAD21 enrichment changes in CTCF^N-Hagfish^ cells showed poor correlation with those in CTCF^N-del^ cells (Pearson’s r = 0.34; Spearman’s ρ = 0.33).

The crystal structure of human RAD21–STAG2 in complex with a CTCF peptide (corresponding to the mouse sequence VSVYDFEEE) revealed the molecular basis of the CTCF–cohesin interaction ^24^. To investigate the molecular basis underlying species-specific differences in the ability of CTCF N-termini to promote RAD21 enrichment, we generated AlphaFold 3 (AF3)-predicted structural models of the N-terminus of CTCF from each species in complex with human RAD21 (residues 281–420) and STAG2 (residues 80–1060). The models suggested that the YxF motif-containing region of mouse, Branchiostoma, and hagfish CTCF could interact with the conserved essential surface (CES) of RAD21–STAG2, whereas the *Drosophila* CTCF N-terminus was not predicted to make this interaction (**Supplemental Figure 3A**; see Methods for details). This difference may explain why the *Drosophila* N-terminus completely fails to support RAD21 enrichment, whereas the *Branchiostoma* N-terminus retains some activity. However, the mouse, hagfish, and *Branchiostoma* N-termini were predicted to engage the CES to a similar extent. Thus, the predicted interaction mediated by the YxF motif alone cannot explain the differences in RAD21 enrichment among these three species. These findings suggest that additional residues outside the YxF motif contribute to robust interactions between cohesin and the CTCF N-terminus. which is predicted to be largely disordered (**Supplemental Figure 3B**).

Together, these results show that the N-termini of invertebrate CTCF proteins have minimal or no capacity to support cohesin recruitment. In contrast, the hagfish CTCF N-terminus can support cohesin enrichment, albeit somewhat less efficiently than its mouse counterpart.

### Co-enrichment of cohesin and CTCF in hagfish tissue

The ability of the hagfish CTCF N-terminus to support cohesin recruitment in mammalian cells prompted us to investigate whether hagfish CTCF performs this function in its native genomic context. We collected inshore hagfish from Kamo Bay, Japan, and mapped genome-wide CTCF and RAD21 occupancy. We identified more than 63,767 CTCF peaks and 29,581 RAD21 peaks in hagfish (**Figure 3A**). The number of RAD21 peaks was comparable to that detected in mouse ESC (50,822 CTCF peaks and 26,055 RAD21 peaks). Although most hagfish RAD21 peaks (61.2 %) overlapped with CTCF peaks, only 28.4% of CTCF peaks overlap with RAD21 peaks, so a substantial fraction was occupied by only one of the two proteins (**Figure 3A**). This overlap was greater in mouse, where 96.1% RAD21 peaks and 49.6% of CTCF peaks overlapped with the other protein peaks.

**Figure 3:**
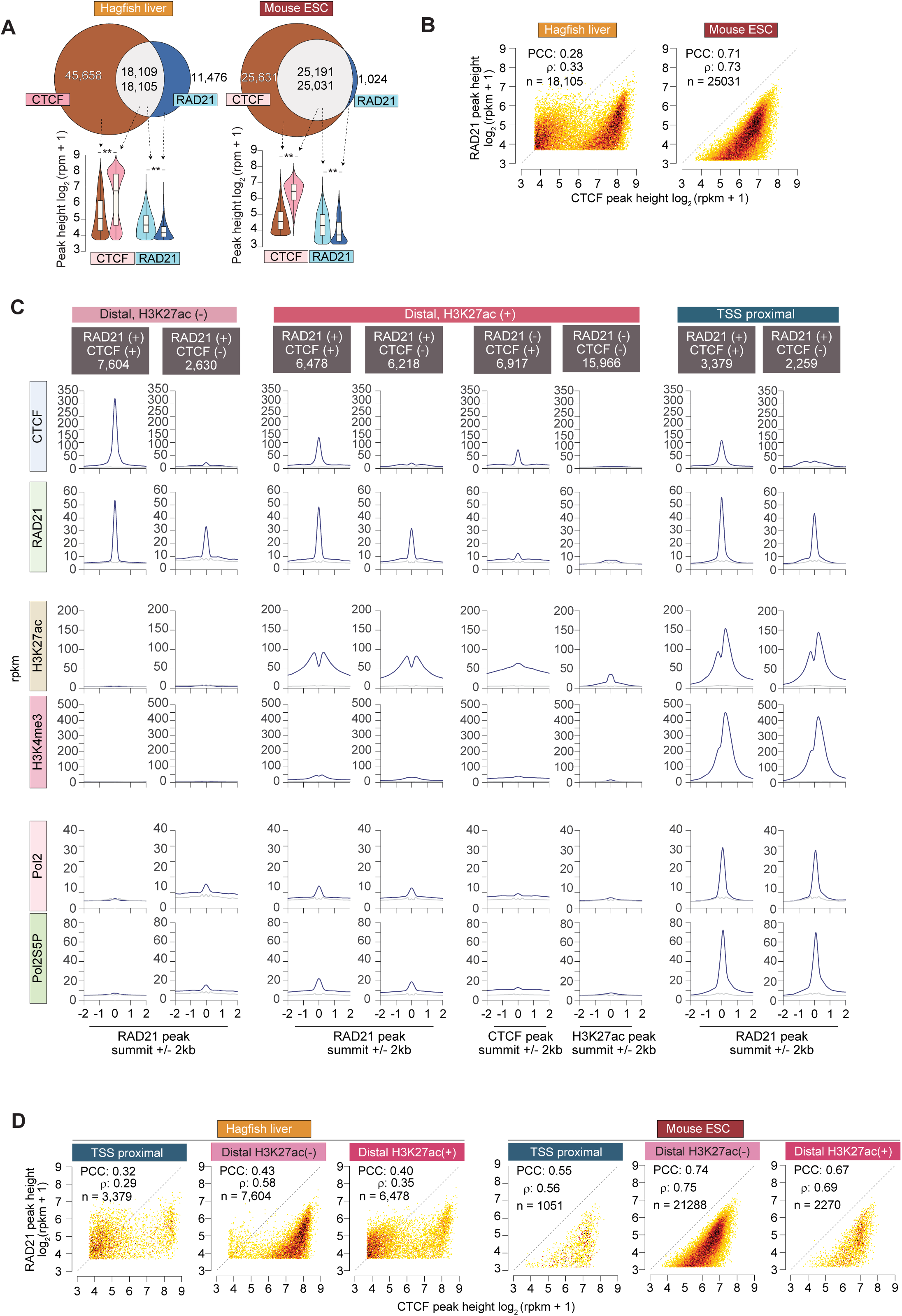
Distinct genomic occupancy of hagfish CTCF and cohesin at promoters, enhancers, and architectural sites. **(A)** Venn diagrams showing the overlap between RAD21 and CTCF ChIP-seq peaks in hagfish liver and mouse ESC. Hagfish ChIP-seq data represent two technical replicates; mouse data were obtained from GSE178982. Violin plots show RAD21 and CTCF peak heights across overlapping and non-overlapping peak sets. In the overlapping region, the upper and lower numbers indicate the numbers of CTCF and RAD21 peaks, respectively. These numbers may differ because a single peak for one protein can overlap with multiple peaks for the other protein, and vice versa. The box plots show the median, upper and lower quartiles; the whiskers show the 1.5× interquartile range. P-values calculated by two-sided Mann–Whitney U-test, followed by correction for multiple comparisons with the Benjamini–Hochberg method. ** *Padj* < 0.001. **(B)** Correlation between CTCF and RAD21 enrichment in hagfish liver and mouse ESC. **(C)** Aggregate ChIP-seq profiles of the indicated histone modifications, Pol II, and Pol II-S5P in hagfish liver. Data represent two technical replicates for CTCF, RAD21, Pol II, and Pol II-S5P, and one replicate for H3K4me3 and H3K27ac. **(D)** Correlation between CTCF and RAD21 enrichment across the indicated genomic regions in hagfish liver and mouse ESC. Regions within ±500 bp of transcription start sites (TSSs) were classified as TSS-proximal, whereas regions located more than 1 kb from a TSS were classified as distal.

Comparison of CTCF and RAD21 enrichment revealed notable differences between hagfish and mouse. In mouse, RAD21 occupancy was strongly correlated with CTCF enrichment (Pearson’s r = 0.71; Spearman’s ρ = 0.73), whereas the correlation was substantially weaker in hagfish (Pearson’s r = 0.28; Spearman’s ρ = 0.33) (**Figure 3B**). This difference reflects two distinct patterns of RAD21 enrichment across hagfish CTCF peaks. At strong CTCF-binding sites, RAD21 enrichment scaled similarly to that observed in mouse. At weak CTCF-binding sites, however, RAD21 enrichment was substantially higher in hagfish than in mouse.

These observations prompted us to examine cohesin occupancy across different classes of cis-regulatory elements. To identify candidate regulatory regions in hagfish, we mapped genome-wide occupancy of several chromatin marks by ChIP-seq (**Supplementary Figure 4**). H3K4me3 enrichment was used to identify candidate promoters, whereas H3K27ac marked both active promoters and candidate enhancers. We also mapped RNA polymerase II (Pol II) and its serine-5-phosphorylated form (Pol II-S5P) to identify transcriptionally active regions. The co-transcriptionally deposited H2BK120ub was mapped as a marker of active gene bodies.

We classified genomic regions as transcription start site (TSS)-proximal (within 500 bp of a TSS) or distal (>1 kb away from a TSS). Distal regions were further classified according to H3K27ac enrichment and then subdivided based on CTCF and RAD21 occupancy. TSS-proximal regions represent candidate promoters, distal H3K27ac-positive (H3K27ac^+^) regions represent candidate active enhancers, and distal H3K27ac-negative (H3K27ac^−^) regions encompass other cis-regulatory elements, including putative architectural CTCF-binding sites.

As expected, H3K27ac and H3K4me3 displayed distinct enrichment patterns at promoters and candidate enhancers (**Figure 3C**). H3K4me3, Pol II, and Pol II Ser5P were strongly enriched at promoters but showed little enrichment in distal regions. In contrast, H3K27ac was enriched at both promoters and candidate enhancers, with characteristic differences in peak shape. At promoters, H3K27ac enrichment was more prominent at the +1 nucleosome than at the −1 nucleosome. At distal regions, H3K27ac enrichment differed between RAD21-positive (RAD21^+^) and RAD21-negative (RAD21^−^) regions. RAD21^+^ regions showed stronger H3K27ac enrichment, with a central RAD21 peak flanked by H3K27ac-enriched regions. In RAD21^−^ regions, H3K27ac enrichment was relatively weaker.

The strongest CTCF enrichment was observed in distal H3K27ac^−^ regions (**Figure 3C**). By contrast, CTCF enrichment was generally weaker at promoters and distal H3K27ac^+^ regions. Notably, where CTCF was detected, RAD21 enrichment was comparable across promoters and both classes of distal regions. Furthermore, many promoter and distal regions showed RAD21 enrichment in the absence of detectable CTCF occupancy.

In hagfish, most CTCF^+^RAD21^+^ peaks were located at promoters and distal H3K27ac^+^ regions (**Figure 3D**). However, the proportion of CTCF^+^RAD21^+^ peaks was higher in distal H3K27ac^−^ regions than in promoters or H3K27ac^+^ distal regions. In contrast, in mouse ESC, only a small fraction of CTCF^+^RAD21^+^ peaks were located at promoters or distal H3K27ac^+^ regions.

### Distinct mechanisms recruit hagfish cohesin in architectural and cis-regulatory regions

To investigate how cohesin is recruited to promoters and candidate enhancers, we examined the co-occupancy of RAD21 and CTCF at TSS-proximal and distal regions.

In hagfish, a large fraction of RAD21 peaks showed no detectable CTCF occupancy (**Figure 4A**). In contrast, nearly all strong RAD21 peaks (top 20,000) in mouse ESCs overlapped with CTCF binding. Even among RAD21^+^CTCF^+^ peaks in hagfish, a large fraction lacked the canonical CTCF-binding motif (**Figure 4B**). This was particularly evident at TSS-proximal regions, where most RAD21^+^CTCF^+^ peaks were CTCF motif-negative. By contrast, the vast majority of RAD21^+^CTCF^+^ peaks in mouse ESCs contained a canonical CTCF-binding motif.

**Figure 4:**
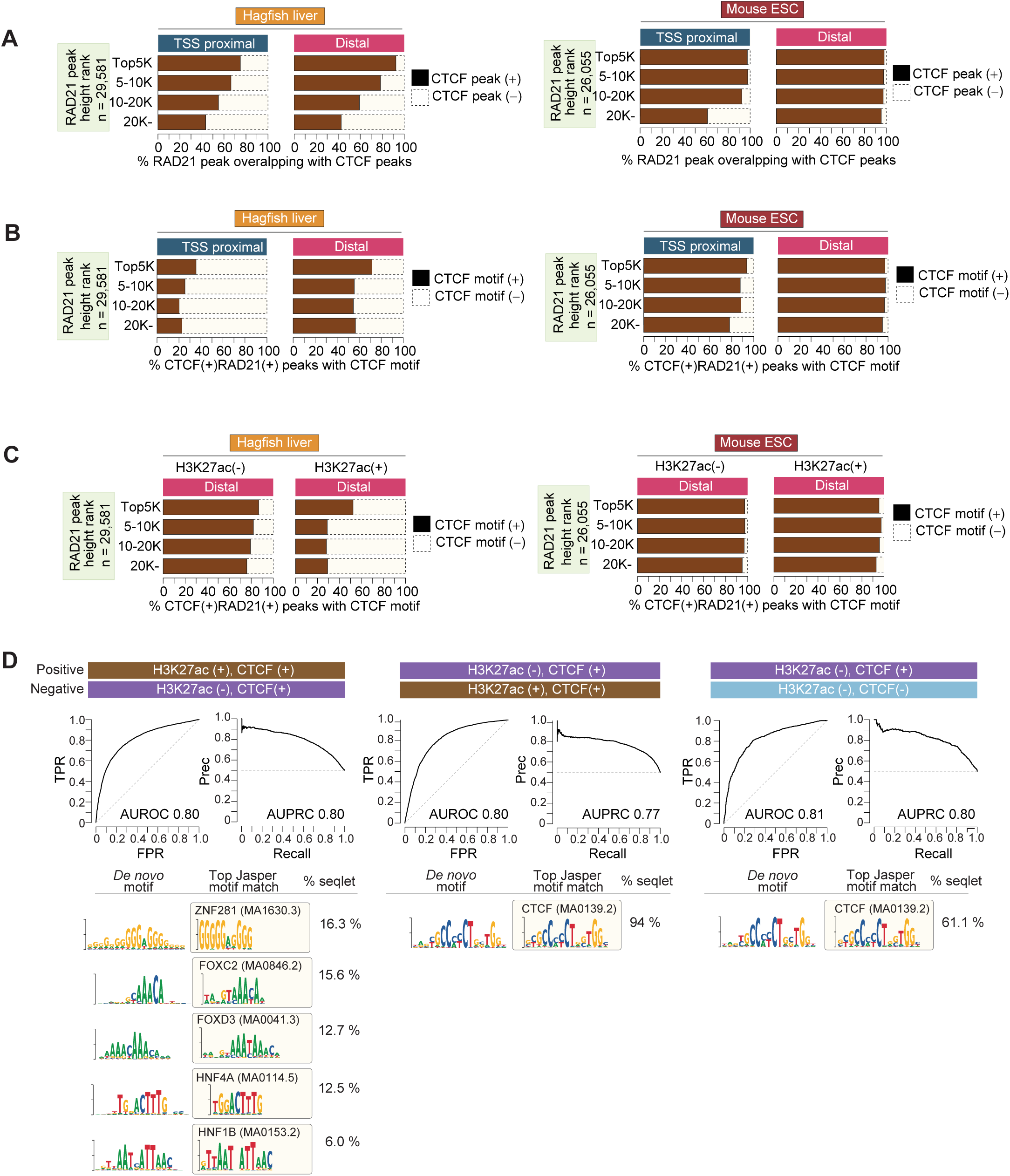
Distinct sequence motifs underlie CTCF enrichment at architectural and non-architectural sites in hagfish. **(A)** Fraction of RAD21 peaks co-occupied by CTCF in hagfish liver and mouse ESCs. RAD21 peaks were stratified by peak height. Shown is the fraction of peaks within each category that overlapped with CTCF binding. **(B)** Fraction of CTCF^+^RAD21^+^ peaks containing a canonical CTCF-binding motif in hagfish liver and mouse ESCs. Peaks were stratified by RAD21 peak height. Shown is the fraction of peaks within each category that overlapping CTCF contained a canonical CTCF motif (Jasper MA0139.2) within its CTCF peak summit +/- 200bp region. **(C)** CTCF motif prevalence among distal CTCF^+^RAD21^+^ peaks in H3K27ac^+^ and H3K27ac^−^ regions in hagfish liver and mouse ESCs. Peaks were first stratified by RAD21 peak height and grouped by H3K27ac status. Shown is the fraction of peaks within each subgroup that contained a canonical CTCF motif. **(D)** Sequence features distinguishing RAD21 peak groups in hagfish liver. Positive and negative peak groups used for training are indicated at at the top of each comparison. ROC and PRC from five-fold cross-validation. Shown is the AUROC and AUPRC values, and de novo motifs from the positive class. Each motif is shown with its best JASPAR match and the percentage of seqlets assigned to it.

Next, we divided distal RAD21^+^CTCF^+^ peaks into two groups based on H3K27ac status. We considered H3K27ac^+^ regions more likely to represent candidate enhancers, whereas H3K27ac^−^ regions were more likely to contain architectural CTCF sites. We then compared canonical CTCF motif (Jasper MA0139.2) prevalence between the two groups. In hagfish, motif prevalence differed markedly: most RAD21^+^CTCF^+^ peaks in distal H3K27ac^−^ regions contained a CTCF motif, whereas the majority of those in H3K27ac^+^ regions did not (**Figure 4C**). In contrast, nearly all distal RAD21^+^CTCF^+^ peaks in mouse ESCs contained a canonical CTCF-binding motif, regardless of H3K27ac status.

The frequent absence of canonical CTCF motifs at regions co-occupied by CTCF and RAD21 prompted us to investigate whether distinct sequence features might underlie their recruitment to these regions. We found that sequence features could distinguish CTCF^+^H3K27ac^+^ regions from CTCF^−^H3K27ac^−^ regions with good accuracy (AUROC = 0.80; **Figure 4D**). This suggests that these two classes of regions have distinct sequence characteristics.

To identify sequence features associated with RAD21 binding in different chromatin contexts, we trained gapped k-mer support vector machine (gkm-SVM)^30^ classifiers on RAD21 peak sequences (±200 bp around peak summit) and extracted de novo motifs from their importance scores using TF-MoDISco. Among distal RAD21^+^CTCF^+^ peaks, the classifier discriminated H3K27ac^+^ from H3K27ac^−^ regions with high accuracy (AUROC = 0.80) (**Figure 4D**). Motifs enriched at H3K27ac^+^ regions matched liver-specific transcription factors, such as ZNF281, FOXA, HNF4 and HNF1, which are key regulators of liver-specific transcriptional programmes^31^. In contrast, H3K27ac^−^ regions were distinguished almost exclusively by the CTCF motif (94% of TF-MoDISco seqlets). Similarly, among distal H3K27ac^−^ regions, RAD21^+^CTCF^+^ peaks were discriminated from CTCF^−^ peaks by the CTCF motif (AUROC = 0.81).

The de novo identified hagfish CTCF motif closely matched the canonical CTCF motif (JASPAR MA0139.2; Tomtom q = 5.8 × 10⁻⁸). This is likely because of the strong conservation of the CTCF zinc fingers between jawless and jawed vertebrates (**Supplementary Figure 5A**). All 20 DNA-contact residues in zinc fingers ZF3–7, which contact the major groove of CTCF-binding sites^32^, are identical between hagfish and human/mouse CTCF (20/20; **Supplementary Figure 5B**). By contrast, 16 of these 20 residues are conserved in *Branchiostoma*, compared with only 11 in *Drosophila*. Both species also show substantial sequence divergence in ZF1–2 (**Supplementary Figure 5C**), which contributes to the full cohesin-retention activity of CTCF^28^. The *Branchiostoma* ZF1-2 show 52-57% identity with human/mouse/hagfish, while *Drosophila* ZF1-2 show 38-52% sequence identity. These differences show that vertebrate and invertebrate CTCF proteins have strongly diverged not only in their N-terminal and ZF1-2 domains. The considerable sequence differences in ZF3-7 imply that vertebrate and invertebrate CTCF may also differ in their DNA-binding sequence preferences.

Together, these results suggest that cohesin is recruited to distal hagfish regions through two distinct modes: interaction with CTCF, as in mammals, or association with tissue-specific transcription factors bound at active enhancers and promoters.

### Hagfish CTCF N-terminus can anchor cohesin loops

To test whether hagfish CTCF acts as a barrier to loop extrusion, we performed Micro-C on hagfish liver. In mammals, CTCF blocks loop-extruding cohesin in an orientation-dependent manner, so that loops form preferentially between CTCF sites whose motifs point toward each other (convergent orientation) ^7,8^. We therefore intersected hagfish CTCF ChIP-seq peaks with the canonical CTCF motif and classified pairs of motif-containing peaks as convergent, divergent or tandem. We then aggregated Micro-C signal around each class (**Figure 5A**). Convergent CTCF pairs showed the characteristic cross of two stripes, one coming from each site, which is the expected sign of orientation-dependent extrusion blocking. Tandem pairs showed a single stripe coming from the inward-facing CTCF site, and divergent pairs showed no stripes. These patterns indicate that hagfish CTCF stalls cohesin in an orientation-dependent manner, as mammalian CTCF does. However, the focal enrichment (“dot”) at the stripe intersection, which marks a stable CTCF-CTCF loop anchored at both ends, was appeared weaker in hagfish compared with mouse. This suggests that hagfish CTCF may be weaker, more permeable barrier to loop-extruding cohesin. This interpretation agrees with our CTCF N-terminus swap experiments in mESCs described above.

**Figure 5:**
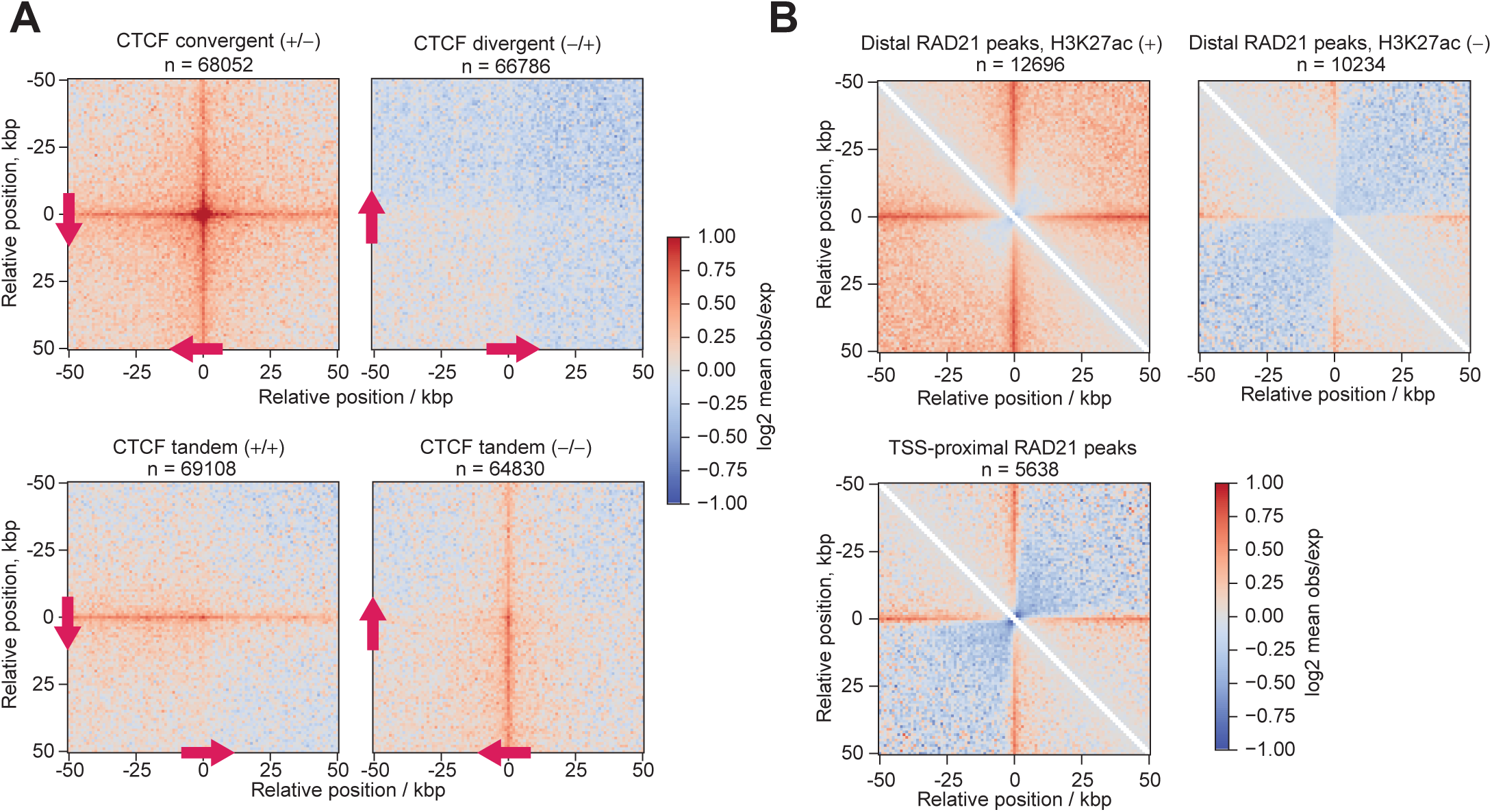
Hagfish CTCF supports cohesin loop anchoring in the native context. **(A)** Aggregate plots of micro-C data at predicted CTCF-CTCF intersections over indicated distances at 1 kbp resolution. Panels from top left to bottom right show convergent, divergent and tandem CTCF orientations with arrows indicating the relative orientation. **(B)** On-diagonal pile-ups at the indicated RAD21-bound TSS-proximal and distal regions at 1 kbp resolution. Distal RAD21-positive regions are further grouped based on H3K27ac status.

To ask whether hagfish cohesin may participate in modulating interactions among cis-regulatory elements, we leveraged our RAD21 ChIP-seq data. We classified RAD21 peaks as either TSS-proximal or TSS-distal, and further subdivided the distal peaks according to H3K27ac enrichment. On the diagonal, pile-up analysis shows that RAD21 peaks that are associated with TSS-proximal regions or distal H3K27ac-positive regions display stripes, whereas the other H3K27ac-negative distal RAD21 peaks show much weaker stripes (**Figure 5B**). This suggest that loop extrusion may serve as a mechanism to promote long-distance interactions among distal cis-regulatory elements in vertebrates.

## Discussion

Chromatin loops are proposed to represent a general feature of animal genomes^3^. Our findings suggest that robust CTCF-anchored loops, by contrast, are likely a vertebrate-specific innovation. We posit that CTCF likely evolved as a bilaterian transcriptional regulator and was later co-opted independently into 3D genome organization in vertebrates and arthropods.

The most distinctive architectural function of CTCF is to halt loop-extruding cohesin^9^, which depends on the YxF motif in its N-terminus^24^. Our findings show that this motif alone is insufficient to confer cohesin-halting activity. Although the motif is conserved in CTCF from *Drosophila*, *Branchiostoma*, and hagfish, their N-termini differ markedly in their ability to support cohesin recruitment. In mouse ESCs, the invertebrate N-termini fail to restore robust cohesin binding, even when fused to mouse CTCF zinc-fingers. Moreover, substantial sequence divergence in ZF1–2, which also appears to contribute to CTCF-dependent cohesin halting^28^, suggests that the cohesin-retention capacity of full-length invertebrate CTCF is likely to be even more limited. In contrast, the hagfish CTCF N-terminus supports clear cohesin enrichment. Consistent with these results, in hagfish tissues ChIP-seq shows cohesin enrichment at distal CTCF-binding sites, and Micro-C identifies chromatin loops. Together, these findings suggest that the ability of CTCF to halt cohesin is a derived property of the vertebrate CTCF. This function likely evolved through sequence divergence in the CTCF N-terminus and may also involve changes in the first two zinc fingers.

These results agree with concurrent preprints reporting TADs and loops in sea lamprey but not in non-vertebrate chordates^33,34^. In lamprey, however, CTCF’s role in anchoring was inferred indirectly from motif enrichment at insulation boundaries. By combining CTCF and RAD21 ChIP-seq with Micro-C, we show directly that, as in mammals, convergently oriented hagfish CTCF sites anchor cohesin loop boundaries.

Despite the shared presence of CTCF-anchored loops, our data reveal two differences between hagfish and jawed vertebrates. First, jawed-vertebrate CTCF appears to halt cohesin more efficiently than hagfish CTCF. Cohesin enrichment is stronger in cells expressing mouse CTCF than in those expressing CTCF with the hagfish N-terminus (**Figure 2**). Second, hagfish cohesin shows a more pronounced enrichment at candidate promoters and enhancers, including sites with weak or undetectable CTCF binding, than is observed in mammals (**Figures 3-4**). Because cohesin also accumulates prominently at promoters and enhancers in *Drosophila* cells ^35^, this feature may not be specific to hagfish.

The presence of CTCF-anchored cohesin loops in both jawless and jawed vertebrates suggests that this function arose in the stem lineage of crown vertebrates, before cyclostomes and gnathostomes diverged. Both groups underwent at least two rounds of whole-genome duplication, but only the first round (1R) is shared^36,37^. Thus, CTCF-anchored loops evolved no later than the first shared genome duplication.

*Drosophila* CTCF also contributes to genome folding, but its mechanism and global importance differ fundamentally from those of vertebrate CTCF. Rather than halting cohesin, *Drosophila* CTCF acts primarily as an insulator-binding protein^5,6^. Moreover, more than a dozen other proteins function as insulator-binding proteins in *Drosophila*^5,6^. CTCF loss weakens only a subset of insulator boundaries and does not globally abolish insulator activity^38^, indicating functional redundancy among insulator proteins. In vertebrates, by contrast, CTCF has a global and largely non-redundant role in cohesin loop anchoring^39,40^, suggesting a more pervasive influence on genome organization and gene regulation. These distinct mechanistic principles support independent co-option of CTCF into 3D genome organization in arthropods and vertebrates.

CTCF is also found outside arthropods and vertebrates, but its architectural role in these other lineages remains doubtful. In many molluscs, the CTCF N-terminus lacks the YxF motif required for interaction with RAD21–STAG2 (**Figure 1**), making cohesin-mediated loop anchoring unlikely. Echinoderms have TADs and retain the YxF motif, but CTCF binding shows no orientation bias at TAD boundaries^41^. Outside arthropods, there is also no evidence for CTCF participating in the formation of insulators. What, then, is the ancestral function of CTCF in other bilaterians?

Beyond its architectural role, mammalian CTCF also acts as a transcriptional activator, binding promoters and regulating hundreds of genes, including housekeeping genes that are ubiquitously essential for cell proliferation^20^. This activity depends on its C-terminal region and is shared with its paralog CTCFL^20^, which cannot anchor cohesin loops^28^. C2H2 zinc-finger proteins are evolutionarily ancient, and transcriptional regulation is one of their conserved functions^42^. We therefore postulate that CTCF originally evolved as a transcriptional regulator, with architectural functions acquired later through lineage-specific co-option. In this model, vertebrate CTCF kept its ancestral activation function and gained the ability to anchor cohesin loops. Arthropod CTCF, in contrast, appears to have lost transcriptional activation function and become integrated into a network of insulator-forming proteins. In each lineage, an ancient transcription factor was repurposed to organize the genome, and in doing so may have supplied the enhancer-promoter specificity on which greater gene regulatory complexity depends.

## Acknowledgments

We thank the members of the Choudhary lab for their helpful discussions. The Novo Nordisk Foundation Center for Protein Research is financially supported by the Novo Nordisk Foundation (NNF14CC0001, NNF24SA0098829). C.C. is supported by the Novo Nordisk Foundation Distinguished Investigator Bioscience and Basic Biomedicine (NNF22OC0074677), and an ERC Advanced Grant (ACT-SIGNAL, 101142708). This work was partially supported by funding from the Institute for Tenure Track Promotion, University of Miyazaki (to Y.H.) and by JSPS KAKENHI Grant Number JP25K02345 (to Y.Y.).

## Author contributions

C.C., H.A, and T.N. conceived the project, E.M., T.B.N.N. contributed to characterization of CRISPR-edited cell lines, Y.Y. and Y.H. obtained hagfish and prepared tissue samples for ChIP-seq and Micro-C analyses; N.K. generated micro-C sequencing libraries and analysed micro-C data, H.A. performed all ChIP-seq analyses, T.N. analysed data and prepared figures, C.C., N.K, Y.H., Y.Y. acquired research funding, C.C., H.A, and T. N. wrote the manuscript writing with input from all co-authors.

## Conflict of interest

The authors declare no conflict of interest.

## Data availability

This project’s sequencing raw data and processed gene tracks are available at the NCBI GEO under accession no. GSE353712. A list of ChIP-seq data generated in this study is provided in **Supplementary Table 1.** Additionally, the following datasets were downloaded and analyzed: gene annotation for mice (GENECODE, https://www.gencodegenes.org), *E. burgeri* (NCBI Genome, https://ftp.ncbi.nlm.nih.gov/genomes/all/GCF/056/783/735/GCF_056783735.1_Ebu_testis_v1/). We reanalyzed the following publicly available sequencing datasets: mouse ESC CTCF (GSE178982), ESC RAD21 (GSE178982), ESC H3K27ac (GSE146328), ESC H3K4me3 (GSE186349), ESC Input (GSE146328), and ESC ATAC-seq (GSE146328).

To review GEO accession GSE353712: Go to https://eur02.safelinks.protection.outlook.com/?url=https%3A%2F%2Fwww.ncbi.nlm.nih.gov%2Fgeo%2Fquery%2Facc.cgi%3Facc%3DGSE353712&data=05%7C02%7Ctakeo.narita%40sund.ku.dk%7C5eadf3dba7f94822c58008df3c2d4b50%7Ca3927f91cda14696af898c9f1ceffa91%7C0%7C0%7C639295837387864273%7CUnknown%7CTWFpbGZsb3d8eyJFbXB0eU1hcGkiOnRydWUsIlYiOiIwLjAuMDAwMCIsIlAiOiJXaW4zMiIsIkFOIjoiTWFpbCIsIldUIjoyfQ%3D%3D%7C0%7C%7C%7C&sdata=260FsWBNjh5funza%2BhZaZ2lqujV%2BjMuoUxWRvHx59tU%3D&reserved=0

Enter token uhexisskxtqjlsn into the box

## Code availability

No new algorithm central to our conclusions was developed in this study. The methods used for the analyses are described in the main text and Methods.

## Materials and methods

### Cell culture

mESCs were purchased from Sigma-Aldrich (E14TG2a; Sigma-Aldrich, cat. no. 08021401). Cells were cultured in a custom-made (C.C.Pro) N2B27 medium consisting of a 1:1 mix of DMEM/F12 and neurobasal medium but lacking arginine and lysine. The medium was supplemented with 1,000 U ml−1 leukemia inhibitory factor (Merck Millipore), 1 µM PD0325901, 3 µM CT-99021 (custom-made by ABCR), 100 µM 2-mercaptoethanol, 0.5 × B27 supplement (Thermo Fisher Scientific), 0.5 × N2 supplement (made in-house or from Thermo Fisher Scientific), L-lysine and L-arginine. The cells were treated with 100-200nM of dTAG-13 and AGB1.

### CRISPR–Cas9 gene editing

The endogenous CTCF locus was edited through the CRISPR–Cas9 system by co-transfecting Cas9-single guide RNA (sgRNA) plasmids with donor homology arms. Full sequences of the homology arms are provided in **Supplementary Table 2**). mESCs were grown in 2i media and donor and sgRNA plasmids were transfected using Lipofectamine 2000 (Thermo Fisher Scientific, cat. no. 11668019), according to the manufacturer’s instructions. After incubating the cells overnight, the transfection mix was exchanged to normal media. Two days after transfection, cells were dissociated using TrypLE (Thermo Fisher Scientific), washed, and seeded onto a new plate and selected using 0.5 μg ml−1 Puromycin (Invitrogen, cat. no. ant-pr-1), 1.75 μl ml−1 hygromycin (Invitrogen, cat. no. ant-hg-1) according to the selection markers used. After six days of selection, the cells were transferred to normal media, and individual colonies were picked and genotyped using KOD Xtreme Hot Start DNA Polymerase (Novagen, cat. no. 71975). Selected cell clones were verified by genome sequencing to confirm genome editing. We first generated a cell line in which the edogenous *Ctcf* locus was modified to generate green fluorescent protein (GFP) and FKBP12^F36V^ (dTAG)-fused mouse CTCF wild-type. In this background, we targeted one of the *Ctcf* allele to generate FLAG-BromoTag (bTAG)-fused, N-terminally deleted mouse CTCF, and bTAG-fused mouse CTCF with N-terminus swapped with the N-terminus of *Drosophila*, *Branchiostoma*, or hagfish. We used microscopy-based imaging to identify clones expressing both GFP-dTAG-fused CTCF wild-type, and N-terminally modified FLAG-bTAG-fused variants.

### Immunofluorescence microscopy

Cells (30,000 cellls/well) were seeded in Geltrex-coated 96-well PhenoPlates (Revvity, #6055300) and allowed to adhere for 24 h. Cells were treated with DMSO, 200 nM dTAG-13, 100 nM AGB1, or a combination of dTAG-13 and AGB1 for 1 h. Following treatment, cells were fixed with 4% formaldehyde (Thermo Scientific, Pierce™, #28908) for 10 min at room temperature (RT) and washed with phosphate-buffered saline (PBS, 3x). Cells were permeabilized with 0.5% Triton X-100 in PBS for 10 min, washed with PBS (3x), and blocked for 1 h at RT with antibody diluent (DMEM with 10% FBS, 0.02% sodium azide, filtered). Cells were incubated with anti-FLAG primary antibody (∼0.4 µg/mL; Sigma-Aldrich, #F3165) overnight at 4 °C. Following washing (PBS, 3x), cells were incubated with Alexa Fluor 647-conjugated secondary antibody (Invitrogen, #A21235) and DAPI for 1.5 h at RT. After three final PBS washes, images were acquired by confocal microscopy (Opera Phoenix) using a 40× objective and 2 × 2 binning.

### Chromatin immunoprecipitation (ChIP) using mouse embryonic stem cells

Mouse embryonic stem cells expressing degron-fused CTCF variants were treated for 4h with DMSO, 100nM BromoTag ® AGB1 (7686, Tocris Bioscience), 200 nM dTAG-13 (6605, Tocris Bioscience), or the combination of AGB1 and dTAG-13. After treatments, cells were cross-linked with 1% formaldehyde (252549, Sigma-aldrich) for 15 min, quenched with 0.2 M glycine for 5 min, washed twice with PBS, then collected by cell lifter, spun down for 2 min at 1,000 g at room temperature. After removing the supernatant, cells were treated with 4 mM disuccinimidyl glutarate (sc-285455, Santa Cruz Biotechnology) in PBS by rotation for 60 min at room temperature, then washed twice with PBS, spun down for 2 min at 1,000 g at room temperature and stored at -80 °C until use. Cells were resuspended in SDS lysis buffer (10 mM Tris-HCl, pH 8.0, 150 mM NaCl, 1% (w/v) SDS, 1 mM EDTA, pH 8.0, cOmplete™ Mini Protease Inhibitor Cocktail (11836145001, Sigma-aldrich)) and sonicated by a Bioruptor Pico (50 cycles of 30 s on and 30 s off, Diagenode), then centrifuged for 10 min at 13,000×g at 20 °C for debris removal. The supernatants were stored at -80 °C until use. To quantify chromatin DNA concentration, DNA was recovered from the supernatant as follows. The supernatant was 3-fold diluted with ChIP elution buffer (50 mM Tris-HCl, pH 8.0, 10 mM EDTA, 1% (w/v) SDS), supplemented with NaCl to 250 mM final concentration, incubated by ThermoMixer C (1100 rpm, Eppendorf) for 20 h at 65 °C to reverse cross-link, then two-fold diluted with water, treated with 0.2 mg/ml RNase A (12091021, Thermo Fisher Scientific) for 30 min at 37 °C, and followed by incubation with 0.4 mg/ml recombinant proteinase K (AM2548, Thermo Fisher Scientific) for 2hr at 50°C. DNA was extracted by phenol-chloroform and precipitated by ethanol with glycogen (R0561, Thermo Fisher Scientific). DNA concentration was measured by Nanodrop (Thermo Fisher Scientific). The size of DNA fragment was analyzed by 4200 TapeStation System (Agilent Technologies). For ChIP, the supernatant equivalent to 100 μg of chromatin DNA was 5-fold diluted with ChIP dilution buffer (20 mM Tris-HCl, pH 8.0, 150 mM NaCl, 1 mM EDTA, 1% Triton X-100, cOmplete Protease Inhibitor Cocktail), incubated with 3 μg of anti-Rad21 antibody (ab992 Abcam) conjugated with Dynabeads M-280 sheep anti-Rabbit IgG (11204D, Thermo Fisher Scientific) or 5 μg of anti-FLAG® M2 antibody (F3165, Sigma-aldrich) conjugated with Dynabeads M-280 sheep anti-Mouse IgG (11202D, Thermo Fisher Scientific) by rotating overnight 4 °C. The immunoprecipitated Protein-DNA complex was washed with ChIP dilution buffer, wash buffer high-salt (20 mM Tris-HCl, pH 8.0, 500 mM NaCl, 2 mM EDTA, 1% (v/v) Triton X-100, 0.1% (w/v) SDS), wash buffer low-salt (10 mM Tris-HCl, pH 8.0, 1 mM EDTA, 250 mM LiCl, 0.5% (w/v) sodium deoxycholate, 0.5% (v/v) NP-40) once respectively, washed with TE three times, then eluted with ChIP elution buffer. The immunoprecipitated DNA was reversed cross-linked and recovered from the eluates as described above.

### Collection of hagfish tissue

The inshore hagfish (Eptatretus burgeri), unsexed and mixed sizes, were caught in Kamo Bay (36°10′35″N, 133°16′44″E; Oki, Shimane, Japan) using eel traps. The fish were then transferred to the laboratory at Shimane University (Matsue, Shimane, Japan) and maintained in an aquarium filled with recirculating artificial seawater (SW; Instant Ocean Premium; Napco, Sumida-ku, Tokyo). The SW was aerated and kept at 15±0.5℃. The aquarium tank was covered by black vinyl sheets to minimize fish stress. On the day of sampling, three individuals (55.2-125.2 g) were randomly chosen, anesthetized, and decapitated before tissue sampling. The SW-diluted eugenol (FA100; Bussan Animal Health, Chuo-ku, Osaka, Japan; final concentration of eugenol was 53.5 mg/L) was used for anesthesia. The care and use of experimental animals complied with Japanese animal welfare laws, guidelines, and policies as approved by the Animal Care and Use Committee of Shimane University (protocol number MA5-01).

### Hagfish tissue ChIP-seq sample preparation

Approximately 250 mg of hagfish liver tissue was trimmed and homogenized in 7 mL of ice-cold PBS supplemented with cOmplete™ Mini Protease Inhibitor Cocktail (Sigma-Aldrich, St. Louis, MO, USA) using a WHEATON® Dounce tissue grinder equipped with a type A pestle (DWK Life Sciences, Millville, NJ, USA). The homogenate was centrifuged at 1,300 rpm for 5 min at 4 °C.

For RAD21 ChIP, the resulting pellet was resuspended in 1 mL of PBS containing 3 mM disuccinimidyl glutarate (DSG; Thermo Fisher Scientific K.K., Tokyo, Japan) and incubated for 30 min at room temperature. The samples were subsequently fixed with formaldehyde at a final concentration of 1% for 10 min. For ChIP targeting CTCF, Pol II, Pol IIS5P, H3K4me3, H3K27ac, and H2BK120ub, crosslinking was performed using 1% formaldehyde alone for 10 min. Crosslinking was quenched by adding glycine to a final concentration of 0.2 M, followed by incubation for 5 min at room temperature. The crosslinked samples were then washed twice with 1 mL of ice-cold PBS supplemented with cOmplete™ Mini Protease Inhibitor Cocktail. The resulting pellets were stored at −80 °C until further use.

ChIP was performed as described above for mouse ESCs, with the following amounts of chromatin DNA and antibodies. For CTCF ChIP, 85–90 µg of chromatin DNA and 10 µL of anti-CTCF antibody (3418, Cell Signaling Technology) were used. This antibody recognizes a C-terminal region of CTCF that is conserved between jawed and jawless vertebrates and has previously been used for ChIP-seq in lamprey^43^. For mapping cohesin occupancy, 75–85 µg of chromatin DNA and 3 µg of anti-RAD21 polyclonal antibody (ab992, Abcam) were used. This antibody was raised against the C-terminal region of human RAD21 (amino acids 571–631), which shares 78% sequence identity with hagfish RAD21. For histone modification profiling, 20 µg of chromatin DNA was used with 5 µL of anti-H3K4me3 antibody (04-745, Millipore), 5 µg of anti-H3K27ac antibody (91193, Active Motif), or 10 µL of anti-H2BK120ub antibody (5546, Cell Signaling Technology). For Pol II and Pol IIS5P profiling, 65–85 µg of chromatin DNA was used with 10 µL of anti-Rpb1 antibody (14958, Cell Signaling Technology) or 10 µL of anti-phospho-Rpb1 CTD Ser5 antibody (13523, Cell Signaling Technology).

### ChIP–seq library preparation

The amount of ChIPed DNA was quantified using the Qubit dsDNA HS Assay Kit (Q32854, Thermo Fisher Scientific). Sequencing libraries were prepared from 1.5–20 ng of ChIP DNA using the NEBNext Ultra II DNA Library Prep Kit for Illumina (E7645L, New England Biolabs), according to the manufactureŕs instructions. Following adapter ligation and uracil excision, the DNA was purified using AMPure XP beads (A63882, Beckman Coulter) and amplified by PCR for 7–11 cycles. The amplified libraries were subsequently purified and size-selected to obtain an average fragment size of 350–400 bp, as determined using a TapeStation system. The pooled libraries were adjusted to 4 nM and sequenced on an Illumina NextSeq 2000 platform.

### Micro-C library preparation

Micro-C was performed as previously described (Author et al., 2026), with minor modifications. 30-40 mg tissue were thwed and incubated in PBS with 0.1% BSA for 20 min, washed and resuspended in MB1 buffer. The suspension was split into five 200 µl aliquots. Each aliquot was digested with MNase (37 °C, 10 min). The digestion was stopped with 1.6 µl EGTA (65 °C, 10 min). The nuclei were pooled and washed twice in 1× T4 DNA ligase buffer, and 10% was kept as an input control. The remaining nuclei went through end-chewing with T4 PNK (37 °C, 15 min) and then Klenow fragment (37 °C, 15 min). Ends were labeled with biotin-dATP/dCTP (25 °C, 45 min), and the reaction was stopped with EDTA (65 °C, 20 min). Proximity ligation was done with T4 DNA ligase for 2.5 h at room temperature. Unligated biotinylated ends were removed with Exonuclease III (37 °C, 15 min). Samples were de-crosslinked overnight at 65 °C with SDS and proteinase K. DNA was extracted with phenol-chloroform and purified with the QIAGEN PCR Purification Kit. The biotinylated fragments were captured on streptavidin beads in B&W buffer and washed with TBW and 0.1× TE. Libraries were prepared on-bead with the NEBNext Ultra II DNA Library Prep Kit for Illumina (end-prep and adapter ligation) following the manufacturer’s instructions. The number of PCR cycles was chosen from a 1 µl test amplification, and the products were cleaned with 0.9× SPRIselect beads. Libraries were quantified by Qubit and TapeStation and sequenced paired-end on an Illumina NextSeq 2000 or NovaSeq X.

### Micro-C data analysis

Micro-C data were processed with distiller (https://github.com/mirnylab/distiller-nf) using default parameters. Pairs were binned into multi-resolution contact maps (.mcool) with cooler at resolutions from 1 kbp to 10 Mb. Maps were balanced by iterative correction (Imakaev et al., 2012), excluding bins with a MADmax value above 30. Technical replicates were pooled with the library_groups option in distiller.

Aggregate peak analysis (APA) and on-diagonal pile-ups were computed with cooltools (v0.6.1), following the cooltools CTCF pile-up tutorial (https://cooltools.readthedocs.io/en/latest/notebooks/pileup_CTCF.html). All analyses used the merged, balanced Micro-C map of both liver replicates at 1 kb resolution (MAPQ ≥ 30). Observed contacts were normalized to cis expected, and the averaged snippets were plotted as log2 mean observed/expected. To define CTCF motif pairs, E. burgeri liver CTCF ChIP-seq peaks were scanned for the JASPAR CTCF motif MA0139.1 within ±100 bp of the peak summit using FIMO (MEME suite v5.5.0; p < 10-4). Motif orientation was assigned from the highest-scoring hit, and peaks with high-scoring hits on both strands were excluded. All cis pairs of motif centres 50-500 kb apart were classified as convergent (+/−), divergent (−/+) or tandem (+/+ and −/−), and APA was computed at the pixel for each pair with ±50 kb flanks.

### Reference genome annotation

Mouse genome annotation and reference genomes were downloaded from GENCODE (GRCm38.p6 release M25, gencode.vM25.basic.annotation.gtf), and *E. burgeri* genome annotation and reference genomes were downloaded from the NCBI Genome portal (Ebu_testis_v1, GCF_056783735.1_Ebu_testis_v1_genomic.gtf). As a reference, the transcript type ‘protein coding’ and ‘lncRNA’ (mouse ESC), and ‘mRNA’ and ‘lncRNA’ (*E. burgeri*) were chosen for representative genes. The transcripts were annotated by the presence of an H3K4me3 ChIP–seq peak (mouse ESC: GSE146328; *E. burgeri* liver: this study) within 1 kb of the TSS. For every gene, isoforms were sorted by the order of H3K4me3 peak height, Pol II enrichment at 1kb around TSS, gene body Pol II enrichment, from highest to lowest, and the top isoforms were chosen as representative transcript.

### Processing of ChIP-seq data

The following datasets were downloaded from GEO and re-processed: mouse ESC CTCF (GSE178982), RAD21 (GSE178982), H3K27ac (GSE146328) and H3K4me3 (GSE186349). Adaptors were trimmed using fastp (version 0.23.4^44^. Reads were aligned to the mouse mm10 or *E. burgeri* Ebu_testis_v1 genome using bwa-mem2 (version 2.2.1, doi:10.1109/IPDPS.2019.00041). Duplicated reads were marked and removed using Picard toolkit tools (version 3.2.0, https://broadinstitute.github.io/picard/).

Peak regions were called by LanceOtron^45^ with the default model (wide-and-deep_jan-2021). Peak height was determined using bamCompare (deepTools v3.5.5)^46^ with the following parameters (centerReads, minMappingQuality 10, 20bp bins, smooth length 400bp, extend reads 200bp, rpm normalization, and subtraction of the input rpm value). The poorly enriched peaks were filtered out using the following thresholds (H3K27ac, H3K4me3, and Pol II, Pol II-S5P, reference mouse RAD21 (GSE178982): peak height > 8 reads mapped per million (rpm), the remaining CTCF and RAD21, peak height > 12 rpm). Broad H3K27ac, H3K4me3, and Pol II, Pol II-S5P peaks proximal within a proximity of 2kb were merged using Bedtools^47^ (v2.31). For the calculation of RAD21 peak height changes after CTCF degradation, RAD21 peaks were called using LanceOtron in each experimental condition for each mutant cell line separately. All the called peaks were combined with the reference RAD21 peaks (GSE178982) and merged if the called peaks were proximal within 200bp proximity. Background-based normalized RAD21 peak heights were analyzed in the reference RAD21 peak (+) regions, and peaks were excluded if the peak height in either condition of comparison was below 12rpm. Background-based scaling factors were calculated from 50Kb tiled genomic regions lacking ATAC peaks and expressed gene body regions, so that the sum of the reads mapped to these background 50kb regions matched between experimental conditions. Peaks were annotated using the gene model described in “Reference genome annotation”, and defined as follows: distal, > 1kb from H3K4me3(+) TSS; TSS-proximal: < 500bp from an H3K4me3(+) TSS. Peaks between 500bp and 1Kb from the TSS were excluded from regions analysis faceted by position relative to the TSS. CTCF and RAD21peaks were defined as overlapping if their peak summit distance is within 200bp.

### DNA Motif analysis

CTCF motifs were identified within +/- 100bp of CTCF peak summits using FIMO^48^ (MEME version 5.5.4) with the JASPAR MA0139.2 matrix (p < 1e-4)^49^. When a window contained multiple matches, the match with the lowest p-value, or if p-values were tied, the match closest to the peak summit, defined the motif position and orientation. To identify sequence features that discriminate RAD21-bound regions of distinct genomic class in *E. burgeri* liver, the Top 20,000 RAD21 peaks were assigned to TSS-distal (> 1kb from an H3K4me3 (+) TSS) classes by H3K27ac and CTCF peak overlap, and +/- 200bp around RAD21 peak summit regions were extracted. For each pairwise comparison, the larger class was randomly down-sampled to the size of the smaller class. Gapped k-mer SVMs were trained with LS-GKM^50^ using the following settings (gapped k-mer, word length 11, match position number threshold 7, up to three mismatches allowed) with 5-fold cross-validation. The ROC and precision-recall curves were computed from pooled-out-of-fold predictions. The GkmExplain^51^ was used to calculate the importance scores of positive sequences from a model trained on all sequences. Seqlets were clustered into de novo motifs with TF-MoDISco-lite (v2.4.0). De novo motifs were compared with JASPAR CORE vertebrate non-redundant PWMs^49^ using Tomtom^52^ (MEME Suite v5.5.9; minimum overlap 5, q < 0.05 or if no motifs are significant, top motif is shown).

### Phylogenetic distribution of CTCF and its N-terminal motifs

CTCF orthologues were collected from per-species protein sets (Ensembl, NCBI RefSeq, FlyBase, UniProt, INSDC and genome-assembly gene models; source per species in **Supplementary Table 3**). Candidate records were required to carry a CTCF-like C2H2 zinc-finger array, detected with the Pfam zf-C2H2 profile (independent E-value < 0.1) against the human reference array (P49711, residues 266–577). Each candidate was assigned to the CTCF family by reciprocal best hit: its best human hit had to be CTCF (P49711) or CTCFL (Q8NI51), and the reverse search within that species had to return the same record. Records whose best human hit was CTCFL (50 records) were classed as CTCFL paralogues and excluded. Truncated gene models were replaced by the matching UniProt record when it was longer, covered the original record (≥95% identity over ≥90% of its length) and spanned at least 290 residues of the zinc-finger array. Two orthologues missed by the automatic search (*Ciona intestinalis* Q4H2I0 and *Ixodes scapularis* A0ACM8DNP7) were added after manual checking. The species tree was built from the NCBI Taxonomy with ETE 3^53^ (version 3.1.3), restricted to Bilateria. The final tree has 527 species. CTCF N-terminal regions (human residues 1–265) were aligned with MAFFT^54^ (version 7.526). Motif columns were anchored on human CTCF (XM_054379399.1): KTYQR at residues 23–27 and YxF at residues 226–228, where the first position was allowed to be Y or F. The tree and annotations were drawn with iTOL^55^ (v6).

### Structure prediction of the CTCF YxF motif and the cohesion CES

Complexes of the CTCF N-terminal region with the cohesin STAG2–RAD21 subcomplex were predicted with AlphaFold 3^56^ (v3.0.3). Each model contained three protein chains: 1. the N-terminal region of CTCF from *M. musculus* (amino acids 1–265), *E. burgeri* (1–423; NCBI ID: XP_085745034), *B. lanceolatum* (1–372; UniProt ID : A0A8K0EUR7) or *D. melanogaster* (1–293; UniProt ID: Q9VS55); 2. *H. sapiens* STAG2 (80–1060); and 3. *H. sapiens* RAD21 (281–420). Multiple sequence alignments were performed separately for each chain using the AlphaFold 3 data pipeline, and structural templates were disabled for all chains. For each of the five seeds, five diffusion samples were generated, and the top-ranked sample per seed (AlphaFold 3 ranking score) was retained. Each model was then superposed onto the crystal structure of human STAG2–RAD21 bound to a N-terminus CTCF peptide^24^ (PDB 6QNX) using STAG2 Cα atoms (80–1048) with gemmi (doi:10.21105/joss.04200, version 0.7.5). The CTCF-binding pocket was defined as the 23 STAG2 and RAD21 residues within 5 Å of CTCF in 6QNX (STAG2 V294, Y297, R298, N325, D326, S327, L329, K330, Y331, G333, W334, K363, L366, F367, S369, R370 and F371, and RAD21 D333, S334, K335, I337, R338 and L341). For the nine-residue motif window aligned to human CTCF 223–231 (VSVYxFEEE; corresponding to *M. musculus* 223–231, *E. burgeri* 380–388, *B. lanceolatum* 332–340, *D. melanogaster* 245–253), we computed the motif Cα RMSD to 6QNX after superposition for the docking assessment (median RMSD: *M. musculus* 2.09 Å, *E. birgeri* 2.11 Å, *B. lanceolatum* 2.84 Å, *D. melanogaster* no motif contact). UCSF ChimeraX^57^ (v1.11.1) was used for the visualization of predicted structures. Overview panels were superposed on STAG2 using the Matchmaker function of ChimeraX, and pocket close-ups were superposed on STAG2 residues 285–380. CTCF residues within 5 Å of the pocket are highlighted.

### Disorder prediction

Per-residue disorder was predicted for full-length CTCF with metapredict (v3.0.1, PID: 34480923). Disordered regions were called with predict_disorder_domains with the following parameter; minimum IDR length 12, minimum folded domain 50 and gap closure 10.

### Statistical analysis and reproducibility

Unless otherwise mentioned, we performed nonparametric statistical tests (two-sided Mann–Whitney U-test) to calculate P-values and corrected for multiple comparisons using the Benjamini–Hochberg method (R package stats v.4.6.2). The Mann–Whitney U-test does not assume normality or equal variances, and we did not formally test these assumptions. No statistical method was used to predetermine sample size. No data were excluded from the analyses except as described in Methods. The experiments were not randomized. The investigators were not blinded to allocation during experiments and outcome assessment.

### Use of AI tools

AI coding assistants (Claude Code, Anthropic; Codex, OpenAI; Grok, xAI) were used to write, test and review scripts for motif analysis and scoring structure predictions, to review human-written analysis scripts, and to assist in checking the Methods text against analysis records. All code was reviewed by models from at least two different vendors and revised until all high-severity issues were resolved. Its outputs were inspected by T.N. No generative AI tools were used to create or alter images; structural figures were rendered in UCSF ChimeraX from the predicted models. Machine-learning analysis tools (AlphaFold 3, LS-GKM, TF-MoDISco) are described in the relevant Methods sections.

## Figure legends

**Supplemental Figure 1:**
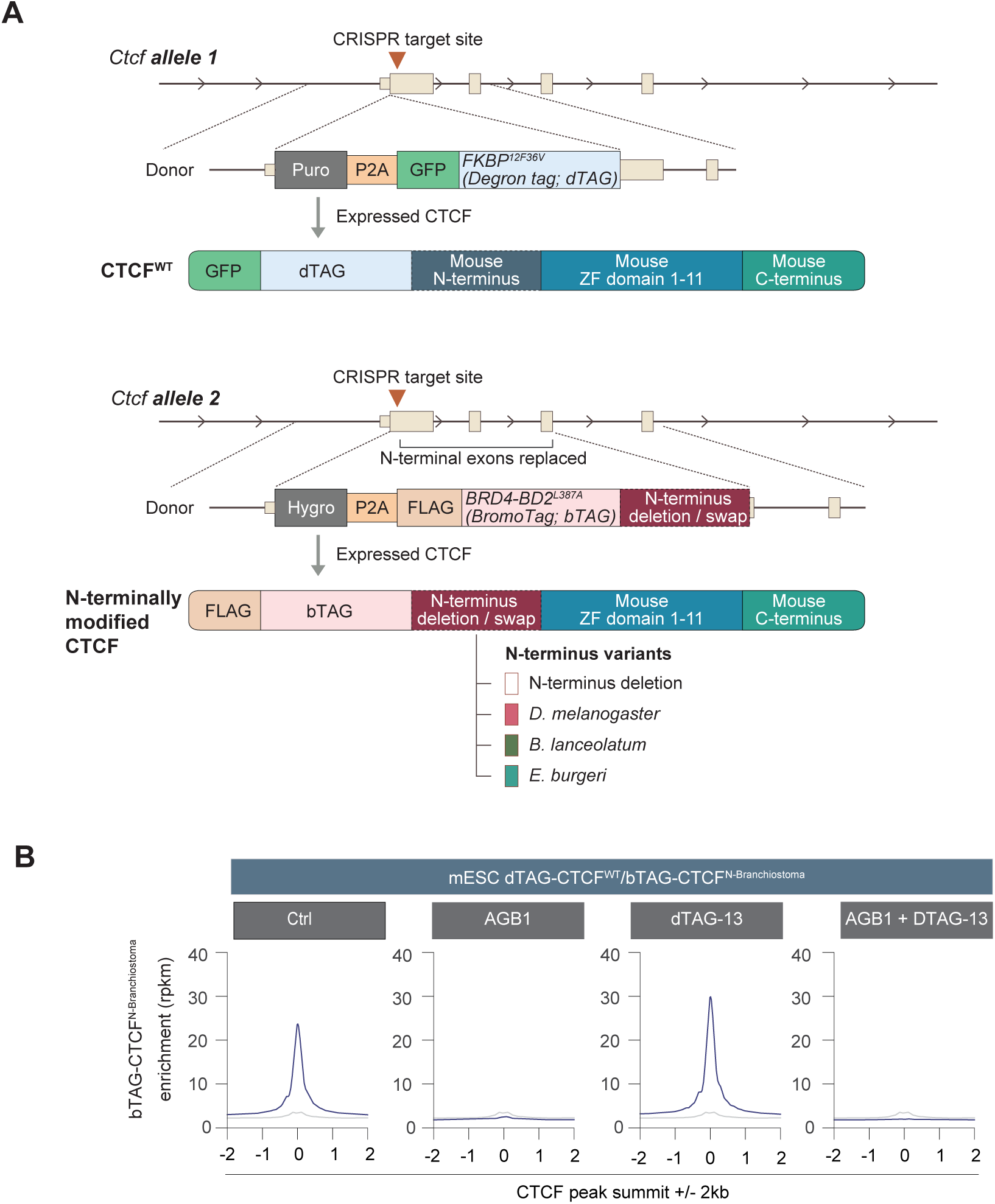
CRISPR-based generation of mouse ESC expressing allele-specific N-terminal CTCF variants, and confirmation of CTCF chromatin binding. **(A)** Schematic of the CRISPR-based strategy used to generate mouse ESC lines expressing allele-specific CTCF variants. In one Ctcf allele, wild-type CTCF (CTCF^WT^) is fused at its N-terminus to GFP and the FKBP12^V36F^ degron (dTAG). In the second allele, CTCF is fused at its N-terminus to a FLAG tag and the BromoTag degron (bTAG). In this allele, the N-terminal region (amino acids 1–265) is either deleted or replaced with the N-terminus from *D. melanogaster* (1–293), *B. lanceolatum* (1–372), or *E. burgeri* (1–423). These modifications generated four mouse ESC lines expressing distinct N-terminally modified CTCF variants. **(B)** Aggregate ChIP-seq profiles showing chromatin binding of CTCF^N-Branchiostoma^. CTCF^WT^;CTCF^N-Branchiostoma^ mouse ESC were treated with dTAG-13, AGB1, or both for 4 h to selectively deplete CTCF^WT^, CTCF^N-Branchiostoma^, or both, respectively. CTCF^N-Branchiostoma^ chromatin binding was assessed by FLAG ChIP-seq. The slight increase in CTCF^N-Branchiostoma^ binding following CTCF^WT^ depletion may reflect reduced competition between the two CTCF variants at shared genomic regions.

**Supplemental Figure 2:**
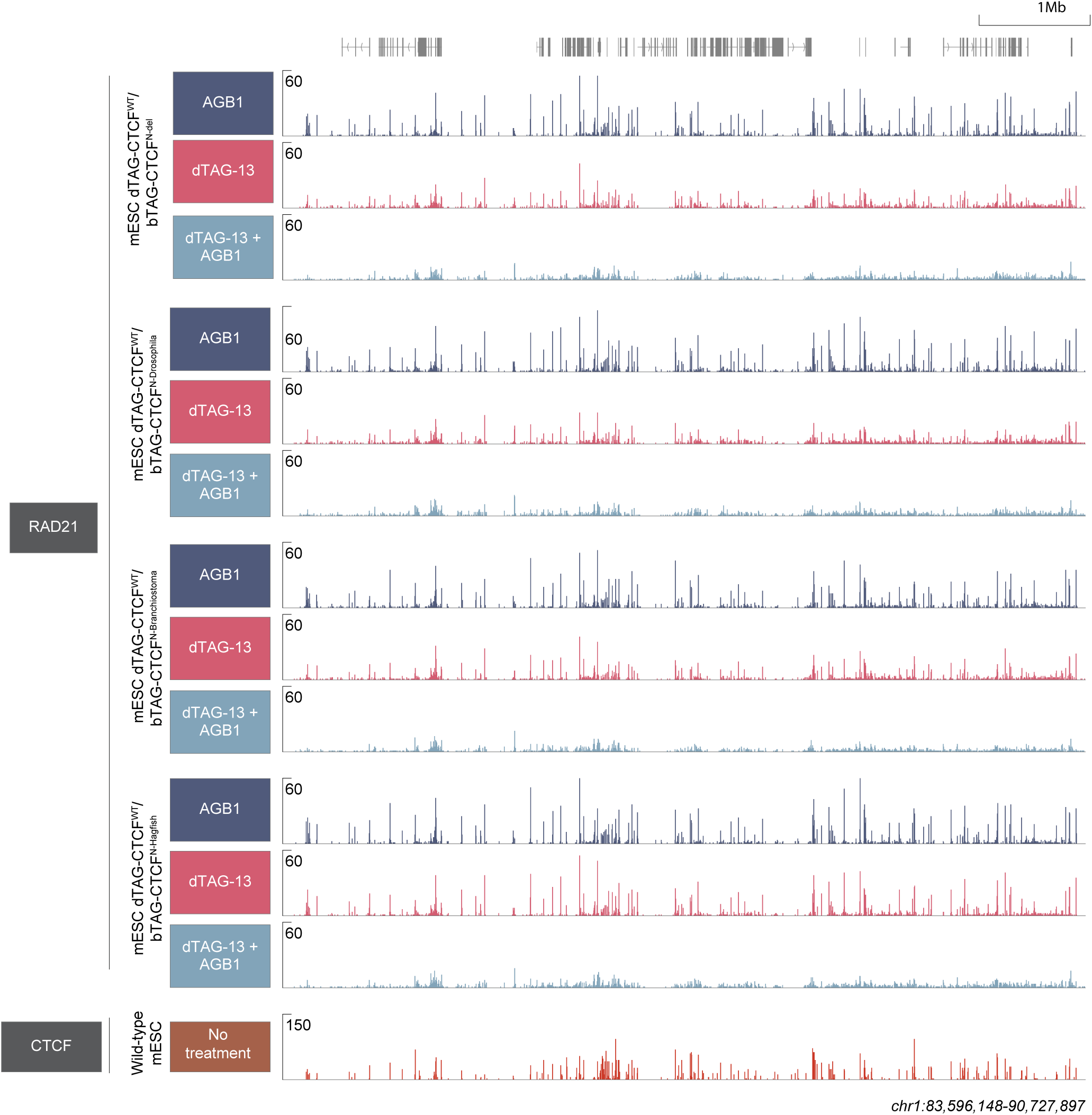
Genome browser views of RAD21 enrichment following selective depletion of CTCFWT or N-terminally modified CTCF variants. The indicated mouse ESC lines were treated with dTAG-13, AGB-1, or both for 4 h, followed by genome-wide mapping of RAD21 occupancy by ChIP-seq (n = 2 biological replicates). dTAG-13 induces degradation of dTAG-fused CTCF^WT^, whereas AGB-1 induces degradation of the bTAG-fused N-terminally modified CTCF variant. Combined treatment depletes both CTCF versions. CTCF ChIP-seq profiles are shown for reference. CTCF ChIP-seq data reprocessed from GSE178982.

**Supplemental Figure 3:**
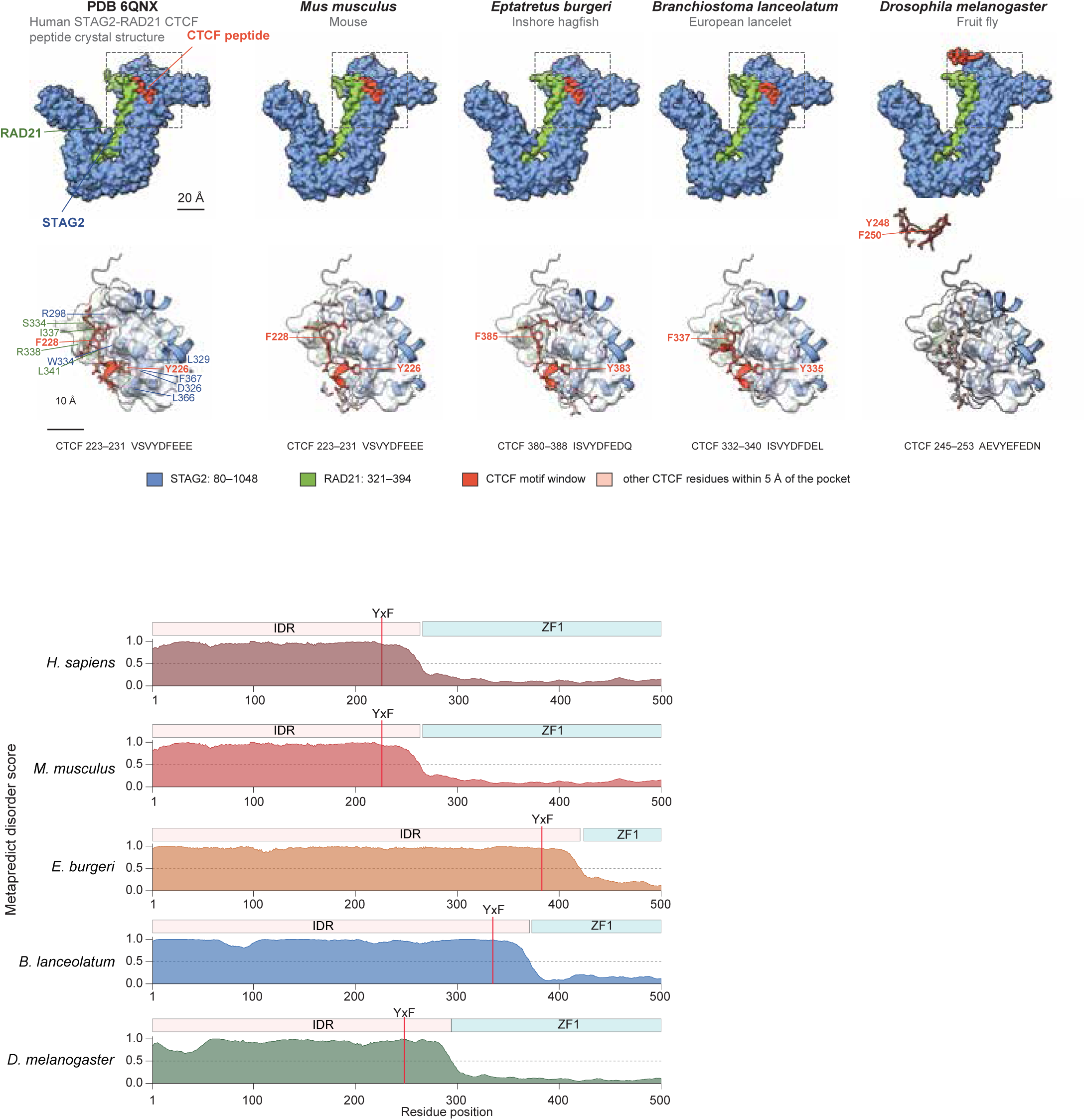
Structural modeling of CTCF N-terminus interaction with human RAD21-STAG2. **(A)** AlphaFold 3 (AF3) models of the CTCF N-terminus bound to human STAG2 (residues 80–1060) and RAD21 (residues 281–420). The left panel shows the crystal structure of human STAG2–RAD21 bound to a human CTCF peptide (PDB 6QNX)^24^. The other panels show AF3 models with the CTCF N-terminus of *M. musculus*, *E. burgeri*, *B. lanceolatum* and *D. melanogaster* with human STAG2 and RAD21. Each complex was modelled with 5 seeds × 5 samples. For each seed, the sample with the highest AF3 ranking score was selected, and of these five, the model with the median motif (corresponding to human VSVYDFEEE) Cα RMSD to 6QNX is shown. For *D. melanogaster*, for which motif Cα RMSD was not computed, the model with the median ranking score is shown. Cohesin is trimmed to the residues resolved in 6QNX (STAG2 80–1048; RAD21 321–394). Top row, surface view. the dashed box marks the CTCF-binding site. Scale bar, 20 Å. Bottom row, close-up of the STAG2–RAD21 pocket: red marks the nine-residue CTCF window aligned to the crystal peptide (human 223–231, VSVYDFEEE), and pale red marks other CTCF residues within 5 Å of the pocket. The Y and F residues of each species’ YxF motif are labelled. Scale bar, 10 Å. **(B)** Predicted disorder score along the CTCF N-terminus of *H. sapiens*, *M. musculus*, *E. burgeri*, *B. lanceolatum* and *D. melanogaster*. The per-residue metapredict disorder score is plotted against Nterminus residues 1–500 of CTCF. Bars mark the predicted disordered region (IDR) and the start of the ZF1. The residue aligned to human CTCF Y226 are marked as YxF. The dotted line marks a disorder score of 0.5.

**Supplemental Figure 4:**
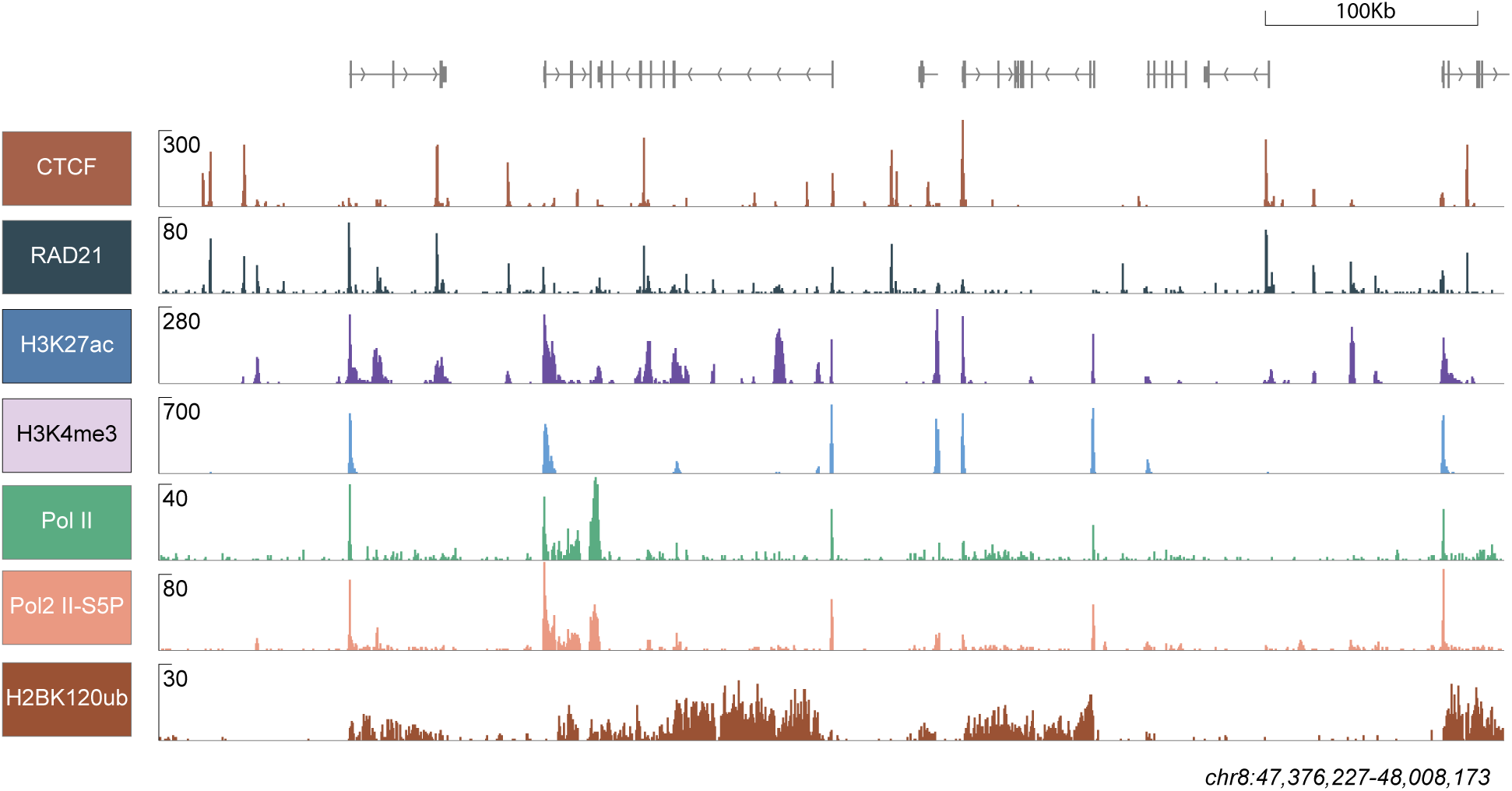
Chromatin mark profiles in hagfish liver tissue. Genome-wide binding of the indicated chromatin marks is profiled using ChIP-seq (n= 2 technical replicates for CTCF, RAD21, Pol II, and Pol II-S5p. n=1 biological replicate for histone marks H3K27ac, H3K4me3, and H2BK120Ub).

**Supplemental Figure 5:**
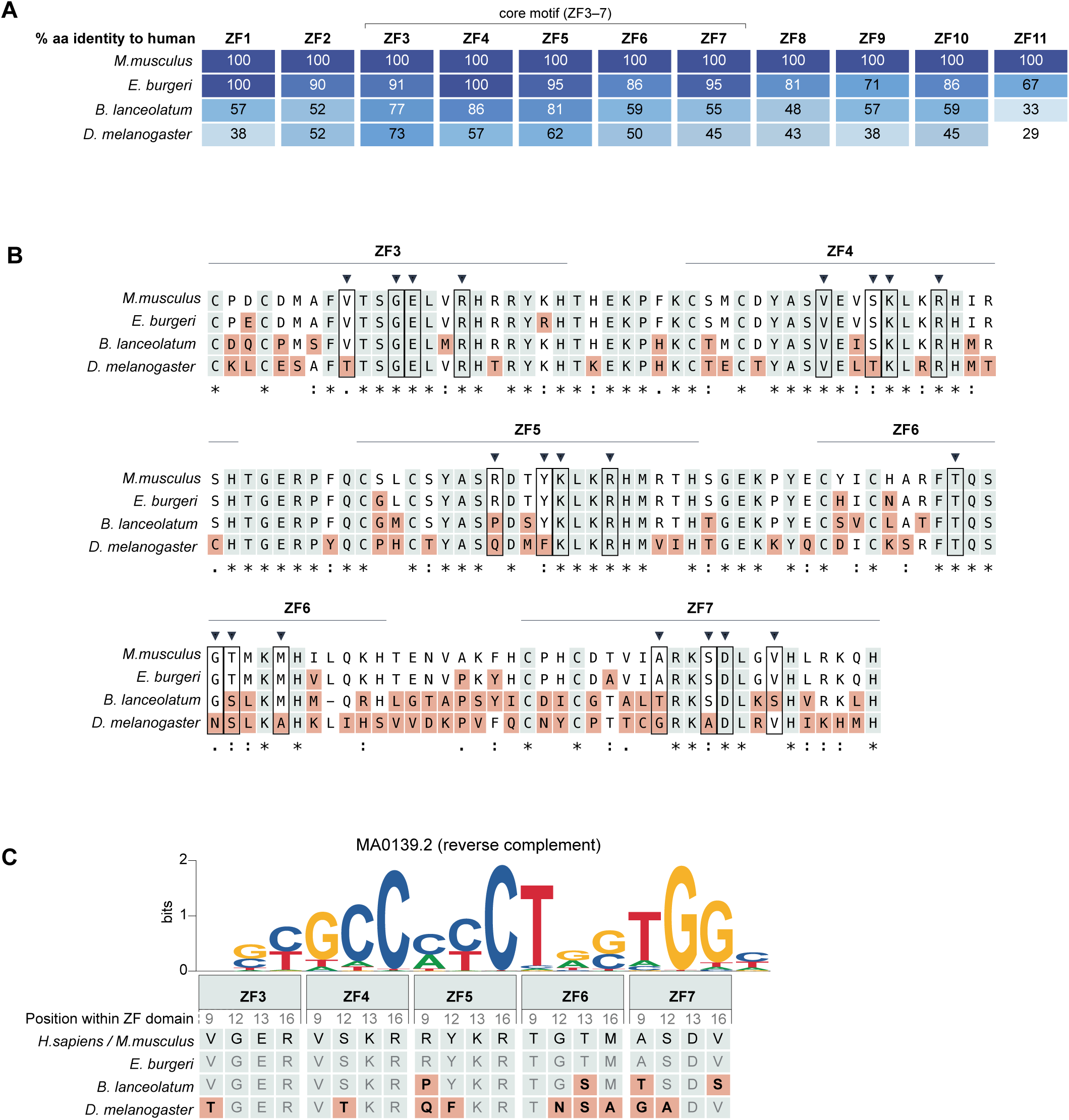
Sequence conservation of CTCF zinc fingers across species. **(A)** Percentage amino acid identity of CTCF zinc fingers 1–11 (ZF1-11) to human CTCF in the indicated species. **(B)** Sequence alignment of CTCF zinc fingers 3–7 (ZF3–7), which mediate DNA binding of the core CTCF motif (MA0139.2). Residues that directly contact DNA and confer sequence specificity are marked with triangles and boxed. **(C)** Amino acid residues in CTCF ZF3–7 involved in DNA binding and their conservation across the indicated species.

## Notes

### Competing Interest Statement

The authors have declared no competing interest.

